# A Neural Model of Schemas and Memory Consolidation

**DOI:** 10.1101/434696

**Authors:** Tiffany Hwu, Jeffrey L. Krichmar

## Abstract

The ability to behave differently according to the situation is essential for survival in a dynamic environment. This requires past experiences to be encoded and retrieved alongside the contextual schemas in which they occurred. The complementary learning systems theory suggests that these schemas are acquired through gradual learning via the neocortex and rapid learning via the hippocampus. However, it has also been shown that new information matching a preexisting schema can bypass the gradual learning process and be acquired rapidly, suggesting that the separation of memories into schemas is useful for flexible learning. While there are theories of the role of schemas in memory consolidation, we lack a full understanding of the mechanisms underlying this function. For this reason, we created a biologically plausible neural network model of schema consolidation that studies several brain areas and their interactions. The model uses a rate-coded multilayer neural network with contrastive Hebbian learning to learn context-specific tasks. Our model suggests that the medial prefrontal cortex supports context-dependent behaviors by learning representations of schemas. Additionally, sparse random connections in the model from the ventral hippocampus to the hidden layers of the network gate neuronal activity depending on their involvement within the current schema, thus separating the representations of new and prior schemas. Contrastive Hebbian learning may function similarly to oscillations in the hippocampus, alternating between clamping and unclamping the output layer of the network to drive learning. Lastly, the model shows the vital role of neuromodulation, as a neuromodulatory area detects the certainty of whether new information is consistent with prior schemas and modulates the speed of memory encoding accordingly. Along with the insights that this model brings to the neurobiology of memory, it further provides a basis for creating context-dependent memories while preventing catastrophic forgetting in artificial neural networks.

## Introduction

Despite the large amount of sensory information in a dynamically changing world, humans develop a structured understanding of the environment, learning to recognize different scenarios and apply the appropriate behaviors. A longstanding goal in neuroscience is to understand how the brain is able to learn these structures quickly and efficiently. The stability-plasticity dilemma asks how the brain is plastic enough to acquire new memories quickly and yet stable enough to recall memories over a lifetime (Abraham & Robins, 2005; Mermillod, Bugaiska, & Bonin, 2013). A related question is how the brain avoids catastrophic forgetting, which is a common problem in connectionist models in which a neural network forgets previously learned skills after being trained on new skills (French, 1999; Kirkpatrick et al., 2017; Soltoggio, Stanley, & Risi, 2017). We believe that the brain is able to avoid catastrophic forgetting and balance stability and plasticity by storing information in schemas, or memory items that are bound together by common contexts. In this paper, we present a neural model that synthesizes existing knowledge of memory processes in the brain and provides a plausible explanation of how schemas are acquired and used in tasks. We also show how the model can be extended to networks used in machine learning applications to avoid catastrophic forgetting in artificial neural networks.

### Brain Areas and Interactions

A prevalent theory of memory consolidation is the Complementary Learning Systems (CLS) theory, which states that memories are first acquired through rapid associations in the hippocampus, and are then gradually stored in long-term connections within the neocortex (Kumaran, Hassabis, & McClelland, 2016; McClelland, McNaughton, & O’Reilly, 1995). The hippocampus replays new memories to the neocortex, interleaving presentations of older memories to prevent catastrophic forgetting. This aligns with hippocampal indexing theory (Teyler & DiScenna, 1986), which states that memories in the form of neocortical activation patterns are rapidly stored as indices in the hippocampus, which are later used to aid recall.

In contrast to CLS, Tse et al. (2007) showed that new memories can be acquired much more quickly if they are consistent with a preexisting schema. In their experiment, rats were trained on a schema for 20 days. The schema layout was a spatial arrangement of six food wells, each containing a different flavor of food. During training, the rats were cued by flavors and released into the arena to find their corresponding wells, which contained food rewards of the same cued flavor. Performance was measured by observing the proportion of time spent digging in the correct well compared to the other wells when cued with a flavor. Performance increased steadily over the 20 days. On the 21st day, two wells were replaced by two new wells with new flavors hidden in them. The rats, when tested on this modified schema, achieved high performance on the memory task despite having had only one day to consolidate. On the 22nd day, some of the rodents received extensive lesions in the hippocampus (HPC). Both conditions of lesioned HPC and control still performed well on the original and modified schema, demonstrating rapid consolidation without hippocampus involvement. The rats were then trained on yet another two wells and flavors within the same schema as before. This was to determine whether the HPC was necessary to learn new information consistent with a prior schema. It was found that the rats with lesioned HPC were unable to learn the new information, thus showing the necessity of the HPC for learning. Further studies by Tse et al. (2011) show activation of plasticity-related genes in the medial prefrontal cortex (mPFC) and related regions, suggesting their involvement in rapid consolidation.

Studies of connectivity between the mPFC and Medial Temporal Lobe (MTL) have yielded theories of how these two areas interact to process schemas. Van Kesteren et al. (2013) provides a framework called SLIMM (Schema-Linked Interactions between Medial prefrontal and Medial temporal regions) to explain how both novelty and familiarity lead to memory encoding. When a sensory stimulus is congruent with a group of preexisting concepts, the mPFC exhibits strong activity and inhibits the MTL, which includes the HPC. If the stimulus is incongruent, the MTL activates. In this way, the two brain areas enhance memory acquisition via different pathways depending on whether the stimulus is familiar or novel relative to prior schemas. While SLIMM provides a general explanation for interactions between novelty and familiarity, it does not include a neural mechanism for how memories are encoded in MTL and mPFC. Eichenbaum (2017) noted that there are direct pathways between the mPFC and MTL, as well as an indirect pathway that routes through the thalamic nucleus reuniens (Re). Eichenbaum hypothesized that theta phase synchronization between the two areas is controlled by the Re to change the directional flow of information when encoding or retrieving. As theta oscillations control many aspects of timing and control in the HPC, they play an important role in schema consolidation.

In addition to the HPC-mPFC pathways for memory consolidation, supporting areas modulate the speed of this consolidation. Kitamura et al. (2017) showed that engrams of contextual fear memories are rapidly formed in the prefrontal cortex via afferent connections from the HPC and basolateral amygdala, suggesting that salience can impact the speed of consolidation. The neuromodulatory system is important for detecting such salience and making quick adaptations (Krichmar, 2008). The basal forebrain (BF) modulates attention and cortical information processing (Baxter & Chiba, 1999), and the locus coeruleus (LC) strongly modulates network activity in response to environmental changes (Aston-Jones & Cohen, 2005). The BF is theorized to represent expected uncertainty when a task is within a known context, whereas the LC signals unexpected uncertainty when context switches occur (Yu & Dayan, 2005). It also drives single trial learning of new information via inputs to the HPC (Wagatsuma et al., 2018). Moreover, the activity from the LC selectively tunes oscillations in the HPC in the theta and gamma range (Berridge & Foote, 1991; Walling, Brown, Milway, Earle, & Harley, 2011), suggesting that this may be one of the mechanisms of single-trial learning.

This type of processing is likely necessary for the type of fast learning that occurs when novel stimuli are introduced within a familiar schema, as seen in Tse et al. Based on this background, we created a neural network architecture with the simulated brain regions and pathways comparable to those described above.

### Contrastive Hebbian Learning

To model hierarchical representations of objects and tasks, a multilayer network can store information of increasing levels of abstraction from input to output layers. Backpropagation is commonly used to train such networks, and has had many successful applications in artificial neural networks (LeCun, Bengio, & Hinton, 2015; Rumelhart, Hinton, & Williams, 1986). However, many view backpropagation as biologically implausible, as there is no widely accepted mechanism in the brain to account for sending error signals backwards along one-way synapses. An alternative account for developing hierarchical representations in the brain may be contrastive Hebbian learning (CHL) (Movellan, 1991), which uses a local Hebbian learning rule and does not require explicit calculations of an error gradient.

Given a multilayered network with layers 0 through L, neuron activations of the kth layer are denoted as vector *x*_*k*_ and weight matrices from the k-1 to kth layer are denoted as *W*_*k*_. Each weight matrix *W*_*k*_ has a feedback matrix of 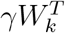 such that every weight has a feedback weight of the same value but scaled by *γ*. The learning process consists of cycling between three phases of the network. The first phase is known as the free phase of the network, in which the input layer *x*_0_ is fixed and the following equation is applied to update neurons in layer *k* from *k* = 1 to *L* at time *t*:

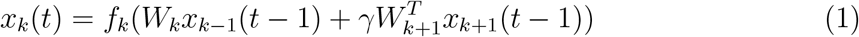

where *f* is any monotonically increasing transfer function. This equation is applied for *T*_*s*_ time steps, which is when network activity converges to a fixed point. The resulting settled activity for *x*_*k*_ is noted as 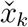, which is the final neural activity for the free phase. The network then transitions to the clamped phase, in which the input layer is fixed as before and the output layer is fixed to the desired target value. Again neuron activities are updated using equation 1 for *T*_*s*_ time steps which allows the network activity to converge. The settled activity for *x*_*k*_ is noted as 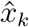, which is the final neural activity for the clamped phase. The third phase combines an anti-Hebbian update rule for neurons in the free phase and Hebbian update rule for neurons in the clamped phase:

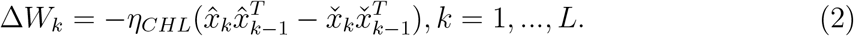

In addition to the added biological plausibility of the local Hebbian learning rule, the repeated alternation between neuron states has been likened to oscillatory patterns in the brain (Baldi & Pineda, 1991), and may serve as a good model for theta synchrony between the hippocampus and neocortex. The fact that preplay and replay of experiences occurs during these times of synchrony suggests that CHL may play a role in rehearsing and reviewing experiences (Diba & Buzsáki, 2007). The scalability of the hierarchical network structure opens the possibility of applying the model to complex tasks. Moreover, as contrastive Hebbian learning was proven to be equivalent to backpropagation for sufficiently small values of *γ* (Xie & Seung, 2003), there is the possibility of expanding the model to problems traditionally addressed by backpropagation. For these reasons of plausibility and scalability, we use a network with contrastive Hebbian learning for our model.

## Methods

Our model consists of two main information streams, the Indexing Stream and the Representation Stream, in a network that is trained on context-dependent tasks such as the one found in Tse et al. (2007). Figure 1 shows an overview of the network.

**Figure 1.**
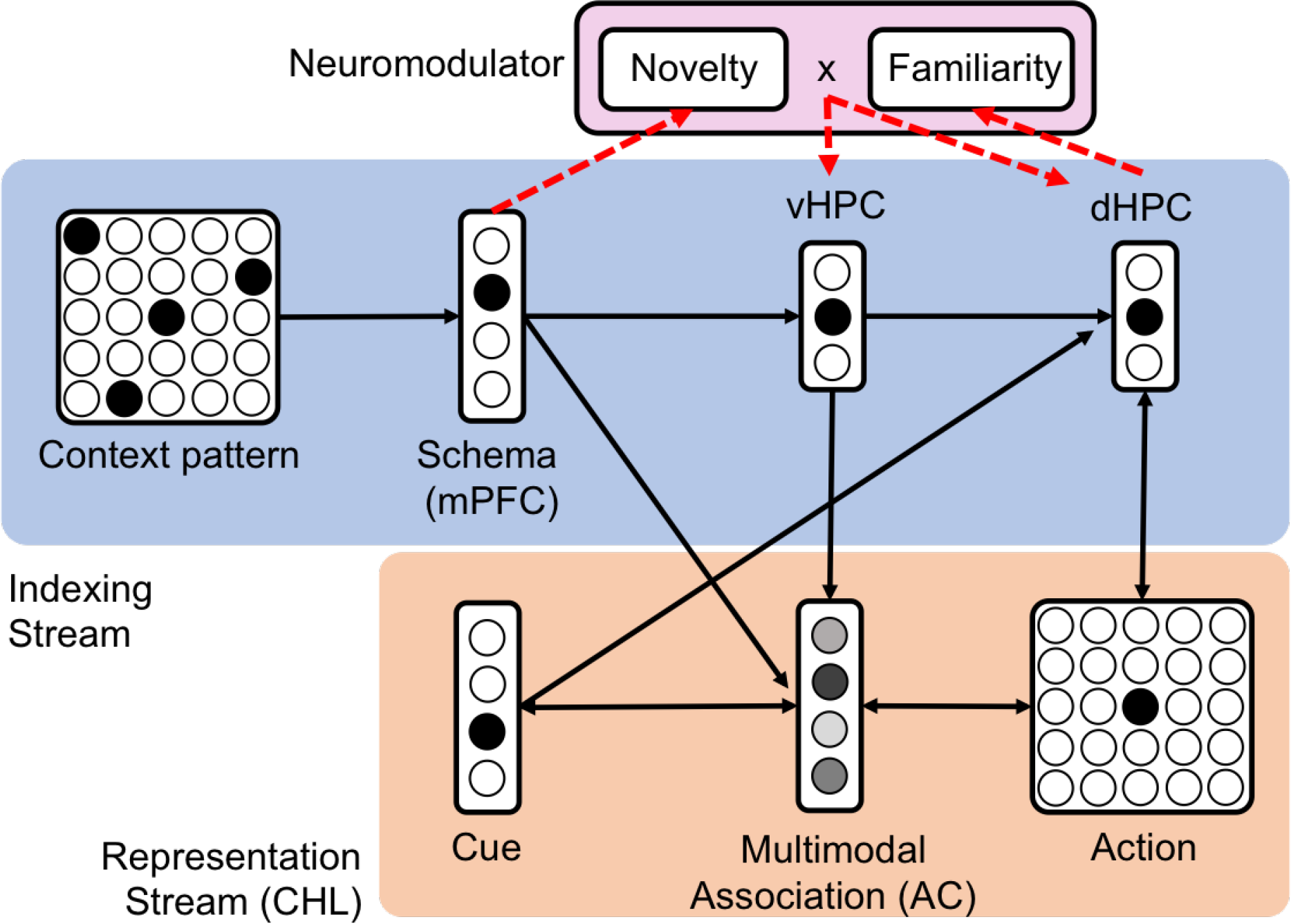
Overview of network. The light blue box contains the areas and projections that make up the Indexing Stream, including the ventral hippocampus (vHPC) and dorsal hippocampus (dHPC). The light orange box contains the areas and projections that make up the Representation Stream of the network. The Representation Stream includes the cue, medial prefrontal cortex (mPFC), multimodal layer (AC), and action layer. Bidirectional weights between layers in the Representational Stream are learned via contrastive Hebbian learning (CHL). Weights from the Indexing Stream are trained using the standard Hebbian learning rule. Dotted lines indicate influences of the neuromodulatory area, which contains submodules of novelty and familiarity. Weights extend to these modules from the mPFC and dHPC. Neuromodulator activity impacts how often the vHPC and dHPC are clamped and unclamped while learning the task via contrastive Hebbian learning (CHL).

### Indexing Stream

The Indexing Stream begins with a context pattern, which can be any encoding of context using patterns of neuron activity. In our case, we used a 2D grid with input activations of 1 if a food well existed in that grid location or 0 otherwise. This input projects to the mPFC for the schema to be learned from sensory input. The dynamic mPFC neuron activity is calculated by the following synaptic input equation at time *t*:

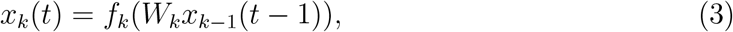

where layer k is the mPFC layer, layer k-1 is the context pattern, and *f*_*k*_ is the Rectified Linear Unit (ReLU) transfer function:

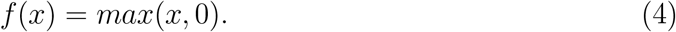

A hard winner-take-all selection is then applied, in which all activations are set to zero except for the one with maximum value. Weights from the context pattern to the mPFC are then trained by the standard Hebbian learning rule:

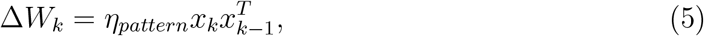

where *x*_*k*_ is the mPFC layer, *η_pattern_* is the learning rate, and *x*_*k*−1_ is the context pattern layer. The weights are normalized such that the norm of weight vectors going to each post-synaptic neuron *i* is 1:

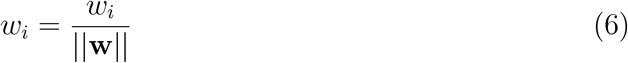

where **w** is the vector of weights going to one postsynaptic neuron, and *w_i_* is an individual weight in **w**. The Indexing Stream continues on to the dorsal hippocampus (dHPC) and ventral hippocampus (vHPC). The dHPC learns an index of mPFC activity using the same synaptic input function and learning rule described in Equations 3-6. The dHPC learns indices in the same way as the vHPC, except that it uses a learning rate of *η*_*indexing*_ and has weights from the vHPC, context pattern, and action selection layer. The term *η*_*indexing*_ is a separate learning rate used for weights that index activity from the Representation Stream. Rather than indexing context, the dHPC indexes triplets of context, cue, and action. This agrees with how context is encoded in the vHPC and specific experiences are encoded in the dHPC in Eichenbaum (2017). It also aligns with the fact that selectivity of encoding increases from the ventral to dorsal end of the HPC (Jung, Wiener, & McNaughton, 1994).

### Representation Stream

The Representation Stream is a multilayered network with sensory cue input areas and the mPFC that encodes the current schema or context. The middle layer of the Representation Stream makes multimodal associations of the sensory input and conjoins context and cue information (AC). While our model only has one middle layer, more layers could be added if the sensory input is complex. The output layer selects an action response to the sensory cue, which in our case is a 2D grid of neurons with an activation of 1 if it is the location of the food cue and 0 otherwise. To train the correct actions, the multilayered network uses the update function and learning rule from contrastive Hebbian learning as described in the background section, with the same transfer function as Equation 4. The alternation of clamped and free phases is controlled by the Indexing Stream. The dHPC alternates between clamping and unclamping the action layer, fixing the action layer at the desired values during the clamped stage of contrastive Hebbian learning. During clamping, the winning neuron in the vHPC gates neurons in the AC. This is done via a static weight matrix of strong inhibitory weights from the vHPC to AC layer, with sparse random excitatory weights that allow only some neurons in the AC to be active. All weights in this matrix are first initialized by a strong negative value of *w*_*inh*_, then a random selection of the weights in this matrix is set to 0. The number of weights selected to be 0 is determined by the value of P, which is a number between 0 and 1 inclusive that defines the proportion of randomly selected weights. This is meant to mimic the effect of hippocampal indexing. While there is little evidence that the HPC projects widespread inhibition to the hierarchical representation areas of the brain, the mix of inhibition and zeroed weights allows specified patterns of neurons in the representations to be active with their usual neuronal activity levels, which is meant to mimic an attentional sharpening effect in the model.

Taken as a whole, the Representation Stream can be viewed as a model of how the neocortex learns hierarchical representations (Hawkins, Ahmad, & Cui, 2017), with the Indexing Stream driving the learning process and preventing catastrophic forgetting by allocating different sets of AC neurons for each task. By using CHL for the Representation Stream, we emulate the oscillatory patterns found in communications between the mPFC and HPC and form an equivalence with backpropagation methods that allows us to expand our model to help improve traditional neural networks in the future.

### Novelty and Schema Familiarity

In the SLIMM framework, the encoding strength is a combination of schema familiarity and cue novelty. Novelty is defined as how infrequently a stimulus has been experienced before, whereas schema familiarity is how frequently a context has been experienced. Furthermore, the SLIMM framework proposes that resonance occurs in the presence of schema familiarity. However, the SLIMM framework suggests that the mPFC inhibits the HPC, whereas we suggest that a combination of schema familiarity and novelty from the mPFC and dHPC, respectively, affect learning by controlling oscillatory activity in the HPC.

In our model, a neuromodulatory area detects novelty in the presence of a familiar schema and modulates the strength of learning that occurs within the Representation Stream. To detect novelty, each neuron in the dHPC projects to the novelty submodule with a starting weight of *w*_*novelty*_. *w*_*novelty*_ represents the baseline level of surprise when a new stimulus is presented. Whenever the activity of the dHPC is updated, the activity of the novelty submodule is found as the rectified weighted sum of inputs from dHPC after applying winner-take-all, as in equations 3 and 4. The weights from the dHPC to the novelty submodule are then updated with an anti-Hebbian learning rule:

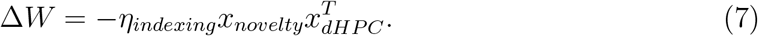

where *W* is the weight matrix of weights from the dHPC to novelty submodule, *η*_*indexing*_ is the learning rate, *x*_*novelty*_ is the activity of the novelty submodule, and *x*_*dHPC*_ is the activity of neurons in the dHPC. The effect is such that the weight between an active dHPC neuron and novelty submodule will experience long term depression and decrease the novelty score of a stimulus after many exposures. Since the dHPC uses winner-take-all, each weight from dHPC represents an individual novelty score for each possible triplet. The activity of the familiarity module is similarly calculated through the weighted sum of inputs from the mPFC after winner-take-all, as in Equation 3. However, rather than a rectified linear unit, we use a shifted sigmoidal transfer function:

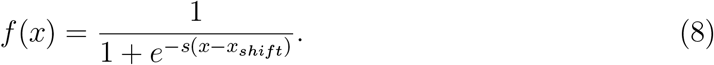

where *−s* is the sigmoidal gain and *x*_*shift*_ is the amount of input shift. The shifted sigmoidal transfer function ensures that the familiarity module requires a baseline amount of training on a schema to be considered familiar with it, and that familiarity does not continue to increase in an unbounded manner with extended exposure. The activity of the familiarity module is thus mostly bimodal, with a very low activity if the schema is unfamiliar and a high activity if the schema is familiar. The effect of the sigmoidal function will be seen in the results. The weights from the mPFC start with the same value of *w*_*fam*_, which is a very small value close to zero that represents low familiarity of schemas prior to training. These weights are updated after mPFC activity is updated, using the Hebbian learning rule from Equation 5 with a learning rate of *η*_*pattern*_. *η*_*pattern*_ is small to model the long-term consolidation of schemas. A final modulation score is then set to the simple product of activity from the familiarity and novelty modules:

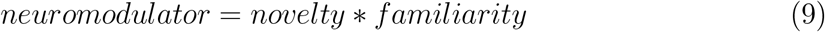

This value determines the number of times the vHPC and dHPC will clamp and unclamp the representation layer in a single trial and thus determine the number of extra epochs in a trial that are added to a default number of epochs, *e*_*default*_:

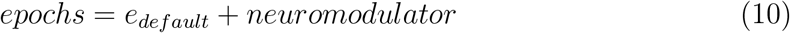

This follows the idea from the SLIMM framework suggesting that resonance occurs in the presence of schema familiarity. However, the SLIMM framework suggests that the mPFC inhibits the HPC, whereas we suggest that information from both areas combines to affect learning.

When training on a task, the network runs trials of training that consist of four phases as shown in Figure 2. Each trial consists of many epochs of training, the number of which is determined by Equation 10. This mimics the high resonance caused by schema familiarity, as postulated in the SLIMM framework. Each epoch consists of an indexing phase, a free phase for CHL, and a clamped phase for CHL. During the indexing phase, the network discovers environmental stimuli and forms indices in the Indexing Stream. The mPFC, vHPC, and dHPC form indices using the unsupervised Hebbian rule and winner-take-all rule described previously. During the Free Phase (Figure 2B) the Representation Stream runs freely, and then during the Clamped Phase (Figure 2C), the the Representation Stream is clamped and the CHL rule is applied. Clamping of the output layer occurs via an input from the dHPC to the action layer using Equation 3. Also during clamping, the AC receives input from the vHPC also using Equation 3, which effectively inhibits most of the AC neurons except for the few that have a weight of 0 from the winning vHPC neuron. At the beginning of a trial, the number of training epochs is undetermined, but tentatively set at *e_settle_* epochs (Figure 2A). During this time, activity levels of the neuromodulator are tracked, and the maximum neuromodulator activity found within this period is used this to determine the ultimate number of training epochs within the trial. After each trial, the performance of the network is measured during the Test Phase (Figure 2D) by presenting a cue to the network and allowing the network to settle on an action. The network parameters used in our network can be found in Table 1.

**Figure 2.**
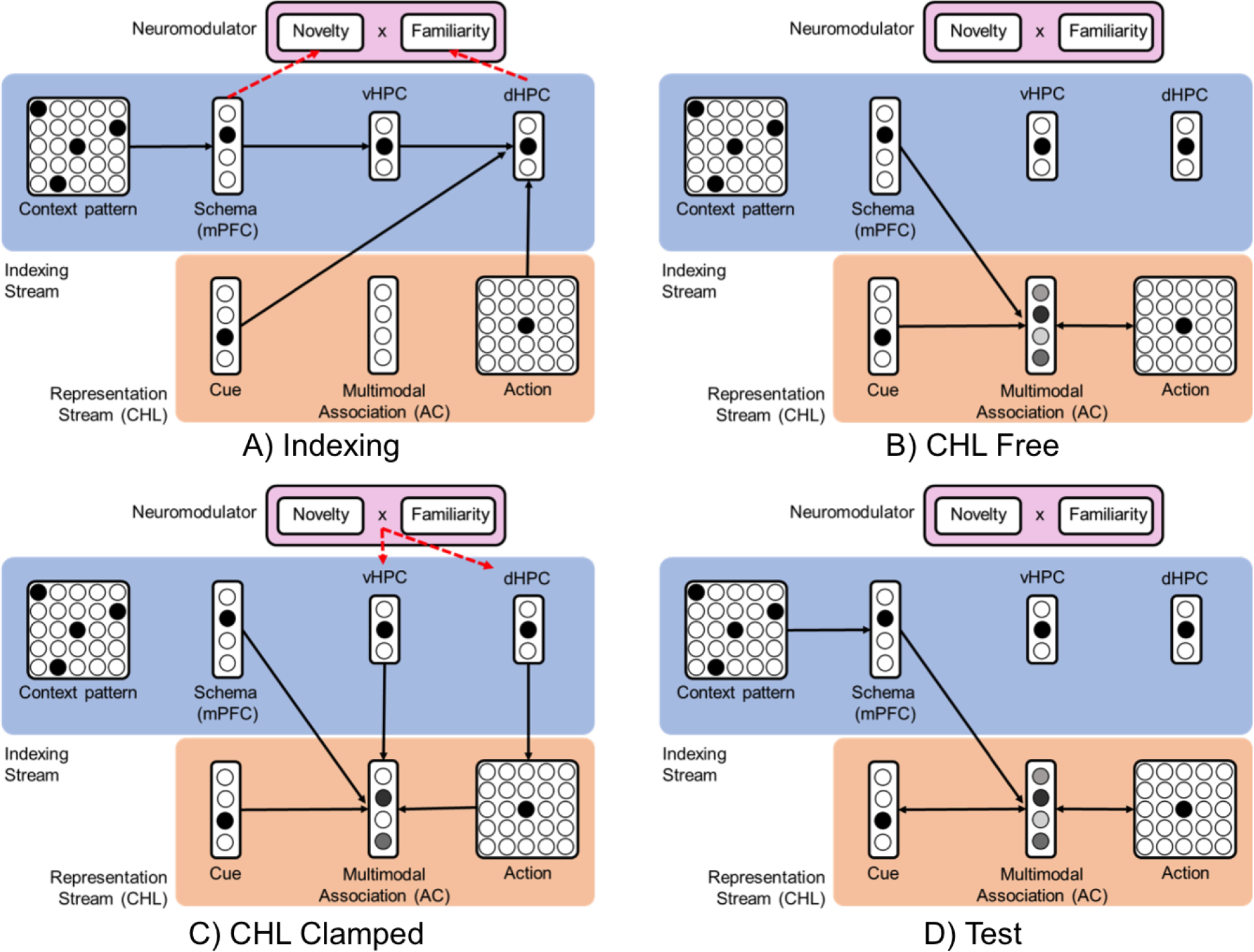
The four phases of a trial of training. A) In the Indexing Phase, the Indexing Stream forms indices of activity. The mPFC learns an index of the context pattern, the vHPC learns an index of the mPFC, and the dHPC learns an index from triples of vHPC, cue, and action. Additionally, the activity of the novelty and familiarity modules of the neuromodulator are calculated, setting the ultimate activity of the neuromodulator to the product of these values. B) The Free Phase of CHL, in which the input is clamped and other CHL areas run free. C) The Clamped Phase of CHL. The clamped value at the action layer is determined by the index in dHPC. The AC layer is gated by the indices in vHPC. The activity of the neuromodulator determines how frequently the clamping occurs within a trial. D) The Test Phase of a network for measuring performance during training and unrewarded probe tests. This is the same as the CHL Free Phase, except that mPFC activity is also calculated.

**Table 1.** Parameters used in experiment.

| Population Sizes |  | Learning Parameters |  |
| --- | --- | --- | --- |
| $N_{cue}$ | 18 | $T_s$ | 5 |
| $N_{mPFC}$ | 10 | $\eta_{indexing}$ | .1 |
| $N_{multimodal}$ | 40 | $\eta_{pattern}$ | .0001 |
| $N_{context}$ | 25 | $\eta_{CHL}$ | .001 |
| $N_{vHPC}$ | 5 | $g$ | .001 |
| $N_{dHPC}$ | 40 | $h_{min}$ | .95 |
| | | $b$ | 8 |
| | | $e_{boost}$ | 1000 |
| | | $e_{default}$ | 600 |
| | | $e_{settle}$ | 20 |
| | | $w_{min}$ | 0.3 |
| | | $w_{max}$ | 0.8 |
| | | $w_{inh}$ | -10 |
| | | $w_{fam}$ | .0001 |
| | | $w_{novelty}$ | 1 |
| | | $s$ | 200 |
| | | $x_{shift}$ | .03 |
| | | $P$ | 0.3 |

### Replication of Tse et Al. Experiments

We validated our model by simulating the experiments performed in Tse et al. (2007). In this case, we simulated a population of 20 rats, each with randomly initialized weights sampled from a uniform distribution in the range of *w*_*min*_ and *w*_*max*_. The arena was discretized into a 5×5 grid, which corresponded to 5×5 grids of neurons for the context pattern and action layers. In our model, each trial consists of multiple epochs to account for replays facilitated by oscillations during sleep and quiet waking periods. In Tse et al. (2007), performance was measured during training by counting the errors in a trial, and non-rewarded probe tests were performed intermittently by recording the amount of time spent searching the correct well when given a food cue. In our model, our test of performance was to present each flavor in a schema during the Test Phase. Upon presenting the cue, the network ran in Free Phase until the activity converged. In the action selection layer, the activity of the neuron corresponding to the correct location of the food was divided by the sum of all of the neurons corresponding to the wells in the arena. This value corresponds to the amount of time a rat would spend digging in a well given a food cue. We used this value to simulate performance during both training and unrewarded probe tests. In the Tse et al. (2007) experiment, the unrewarded probe test was conducted with a food cue and no food reward in the correct well, thus limiting learning for those trials. Although our experiment does not model reward representation in the brain, we approximate unrewarded probe tests by not updating weights after cueing and running the network.

A timeline of the three replicated experiments can be seen in Figure 3. The first experiment was to train the network on the Schema A layout for 20 trials (Figure 3A; PTs 1-3). On the 21st trial, two of the PAs were replaced by two new ones (Figure 3A). The original schema with two new PAs was trained for one trial, and then a probe test was performed (Figure 3A; PT 4). After that, the network was split into a control network and a lesioned HPC group. The HPC group was cloned with weights from the original network and had all connections to and from the vHPC and dHPC removed. Another probe test was performed to see whether both groups could still recall the original schema as well as the schema with two new PAs (Figure 3A; PT 5). Next, the two new PAs were replaced with yet another two new PAs and trained on both groups for one trial. Another probe test was performed after this (Figure 3A; PT 6). The second experiment was performed on the resulting networks from the the first experiment (Figure 3B). For 16 trials, both conditions were trained on an entirely new schema, Schema B, and a probe test was conducted (Figure 3B; PT 7). After this, the groups were retrained on the original schema for 7 days, with yet another probe test at the end (Figure 3B; PT8). For the third experiment, a newly instantiated network was trained on two schemas, one consistent and one inconsistent. We used Schema A for the consistent condition (Figure 3C). For the inconsistent condition, we trained on Schema B, except with the locations of the flavors scrambled on every third trial. The network was trained in an alternating schedule between consistent and inconsistent schemas. Probe tests were performed for the consistent and inconsistent conditions. After this, two PAs from both the consistent and inconsistent schemas were replaced by two new PAs and trained on the rats, and another probe test was conducted.

**Figure 3.**
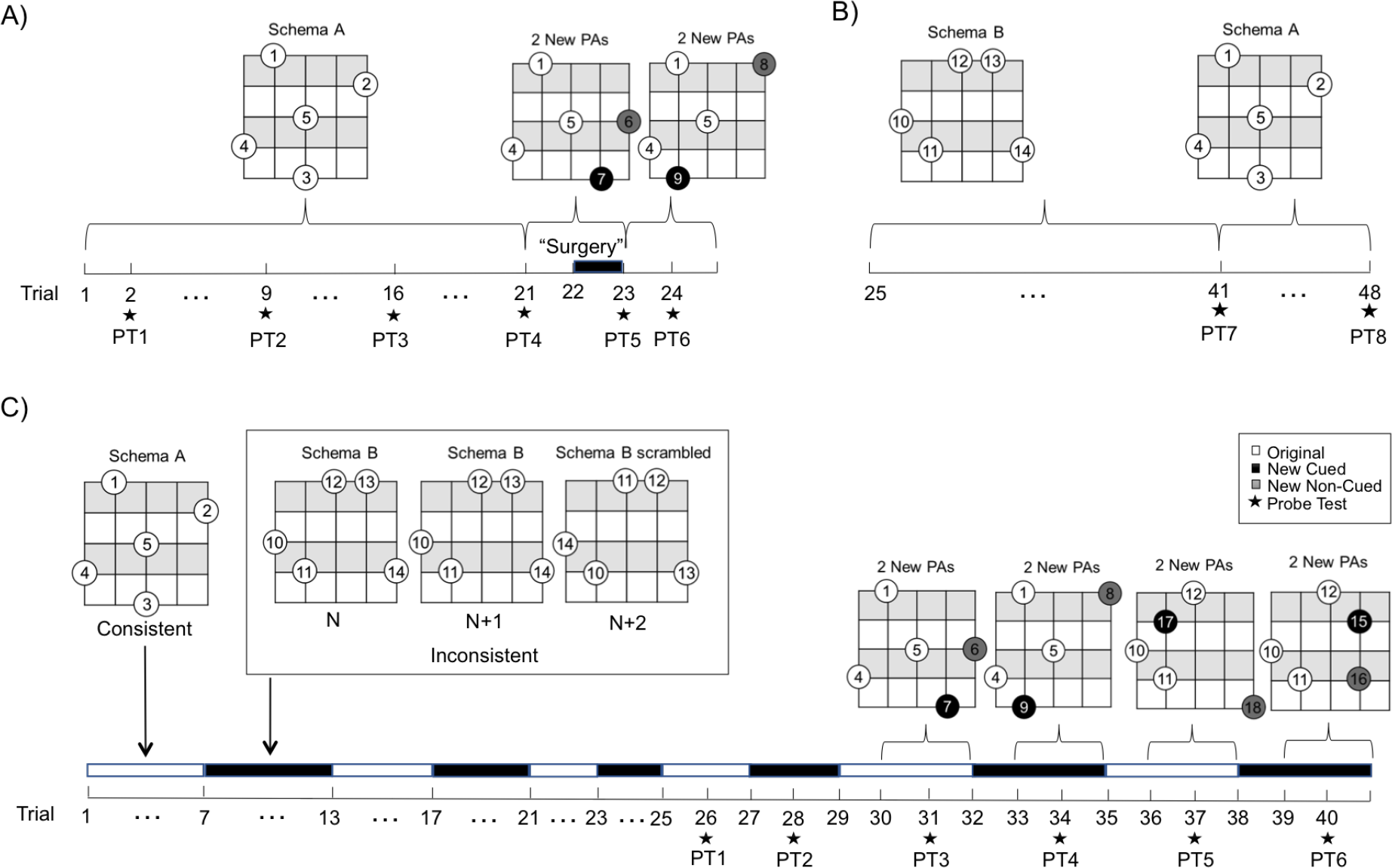
Timeline of experiments. A) Experiment 1 timeline. Schema A is trained for 20 days. Two new paired associations (PAs) are introduced on day 21. Lesioning occurs on day 22, and yet another two PAs are introduced on day 23. Probe tests are performed throughout. B) Experiment 2 timeline. Schema B is trained for 16 days, then Schema A is retrained for 7 days. C) Experiment 3 timeline. Training alternates between consistent and inconsistent schemas. For inconsistent schemas, every 3rd trial has a scrambled version of the schema. White bars represent a consistent schema, in which Schema A is presented every time. Black bars represent an inconsistent schema, where every third trial is presented a scrambled version of Schema B.

## Results

### Experiment 1

The goal of the first experiment was to show that new information matching a schema can be quickly learned, and that the HPC is necessary for this learning. Figure 4A shows that the model was able to gradually learn Schema A over 20 trials of training. Figure 4B confirms this with probe tests, which show the proportion of neuron activity corresponding to the correct action given a cue. Figure 4C shows the probe test results after training for one day on Schema A with two new PAs added. During the Test Phase, the higher dig time in the correct well shows that the new PAs were learned in just one trial. Then in Figures 4D and 4E, upon splitting into two conditions of lesioned HPC and control, both groups were still able to recall the original schema and the two new PAs. This suggests that the new information had been consolidated within a short period and no longer required the HPC for retrieval. We believe this was due in part to the activity of the neuromodulator, detecting when new information was present within a familiar schema and increasing the consolidation accordingly. When training yet another two PAs on both conditions in Figure 4F, the group with the lesioned HPC was unable to learn at all. This was because the CHL mechanism was unable to update the weights properly, as the HPC was responsible for driving the index-based clamping and unclamping of the output layer.

**Figure 4.**
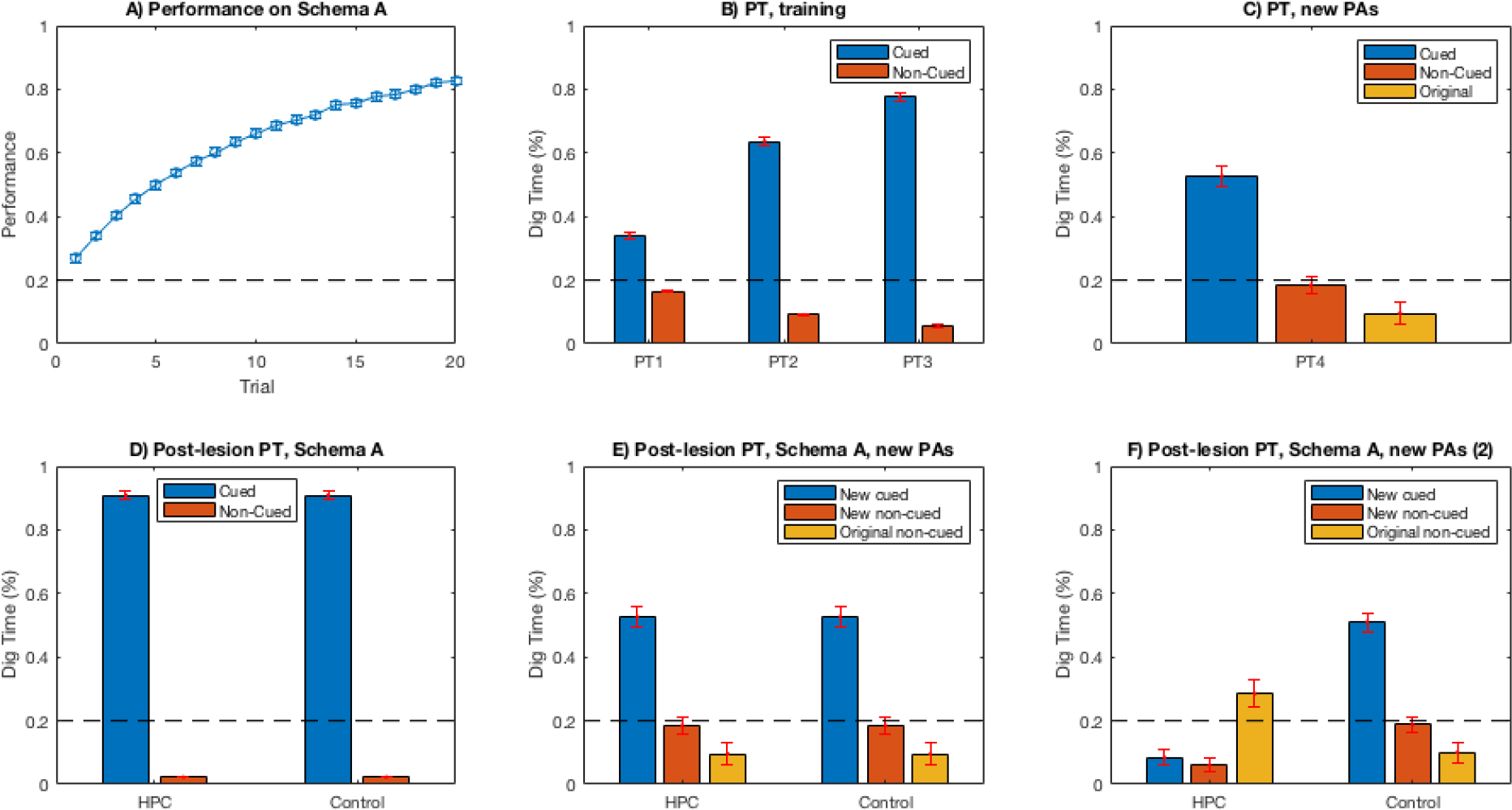
Results of replicating the first experiment of Tse et al. A) The performance over 20 trials, showing a gradual increase. B) Probe tests of trials 2, 9, 4, and 16, showing the proportion of activity of the correct well given a food cue compared to activity of the incorrect wells. C) Probe test after training the new PAs for 1 day, in which one of the new pairs is cued and the activities of the correct well, well of the other new pair, and original Schema A wells are compared. The new PAs were learned within one trial, as the dig time for the cued pair was significantly higher than the rest. D) Probe tests of Schema A after splitting into HPC-lesioned and control groups. Both conditions retained knowledge of the schema. E) Probe tests of the new PAs after splitting into HPC-lesioned and control groups. Both groups recalled the new PAs equally. F) Probe tests of Schema A after training another two new PAs. The HPC group could not learn.

Figure 5 shows the weights of the network after training on the first experiment. Weights from the context pattern to mPFC show the development of a distinct schema pattern encoded with stronger weights where the wells are located. Weights from the mPFC to the AC show the effect of the gating, in which the mPFC neuron representing Schema A is associated with a set of neurons in the AC. Weights from the AC to action layer show that the actions are dependent on activity from select AC neurons, suggesting that the AC neurons are learning specific features useful for action selection. Weights going from the vHPC, cue, and action to the dHPC are displayed in one matrix to show clear encodings of triplets with one neuron from each of the three areas. Weights from mPFC to vHPC show that there is not necessarily a one-to-one mapping from a winning mPFC neuron to a vHPC neuron, but that the schema information is transferred in a distributed manner.

**Figure 5.**
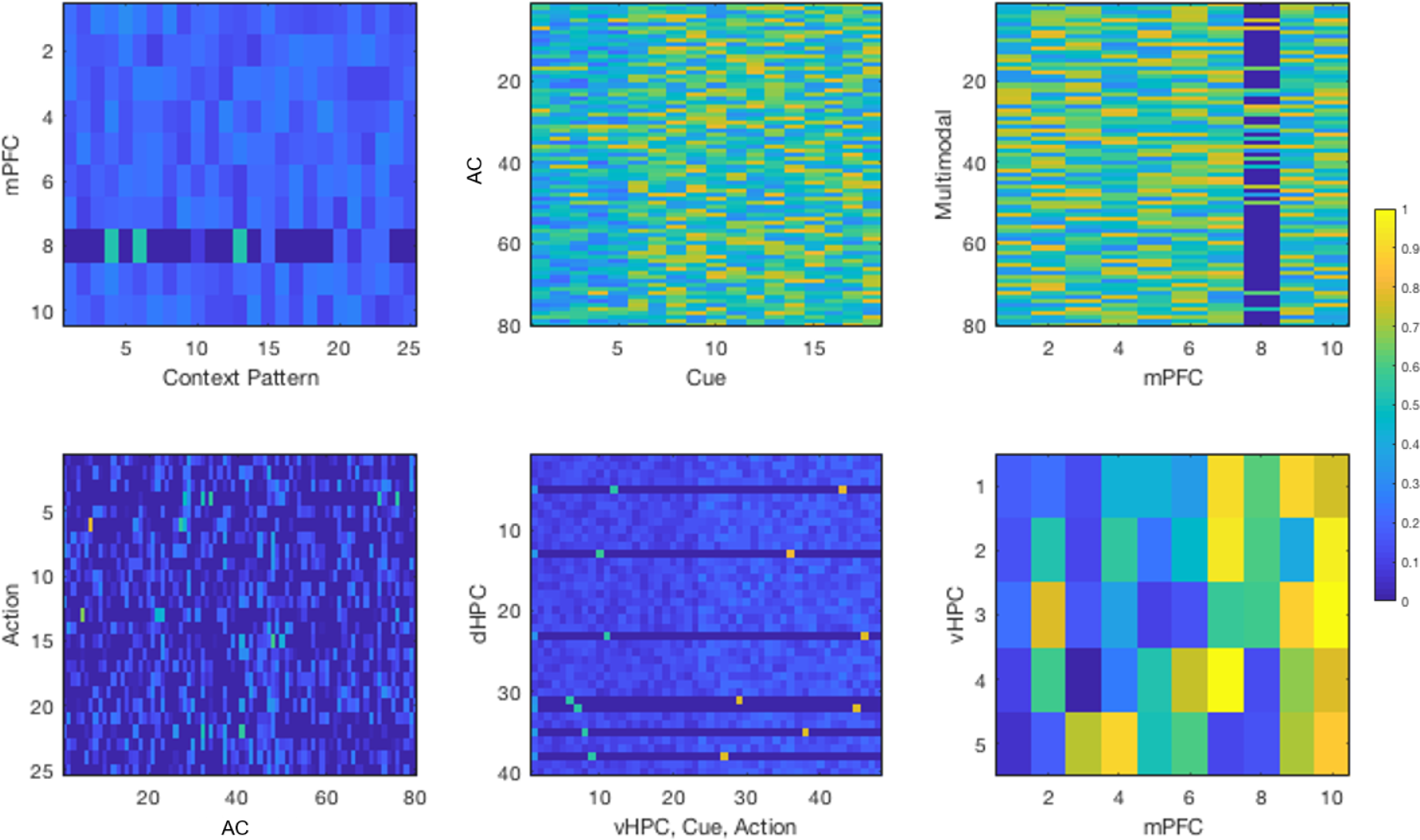
Weight matrices of the network of one simulated rat after simulating all of the first experiment. Rows represent post-synaptic layers and columns represent presynaptic layers.

Our results for the first experiment were able to capture most of the effects seen in Tse et al. (2007). We were able to show that information is acquired rapidly when consistent with a prior schema and that the HPC is necessary for acquisition.

### Experiment 2

The purpose of the second experiment was to test whether multiple schemas could be learned by the same network, and whether the HPC was necessary for this. Figure 6A shows that the control group was able to learn Schema B quickly. HPC was unable to learn, staying at chance levels the entire time. This is shown again in the probe tests in Figure 6B. When the HPC-lesioned model was retrained on Schema A in Figure 6C, it had good performance the entire time and retained the information learned prior to surgery, but did not increase. The control group displayed a minute decrease in performance of Schema A at the beginning, but quickly regained prior performance. The decrease was likely due to small overlaps in the gating patterns from vHPC. This is confirmed again in the probe tests in Figure 6D.

**Figure 6.**
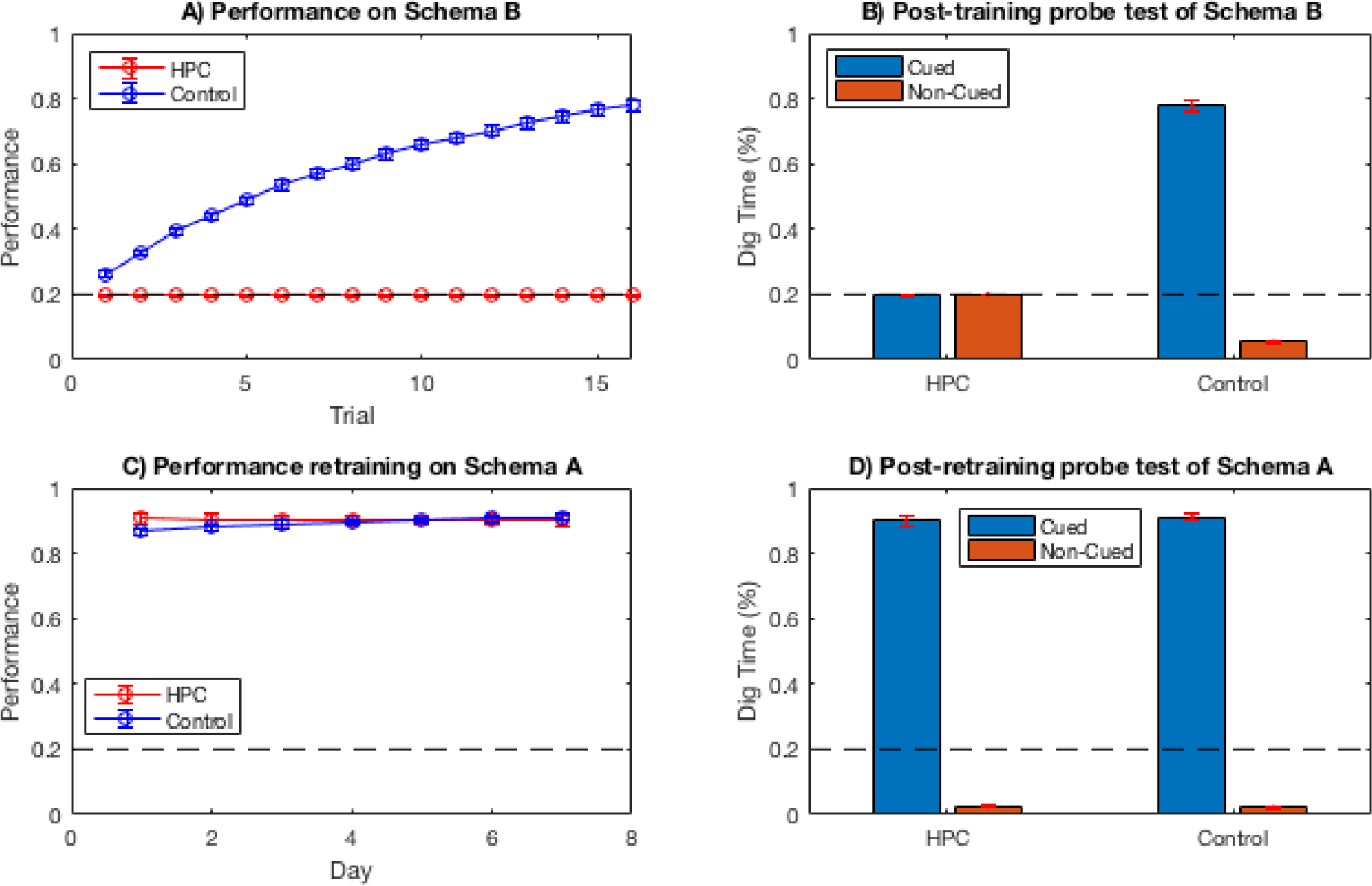
Results of replicating the second experiment of Tse et al. A) The performance over 16 trials of training Schema B. The control group was able to learn the new schema while the HPC group was not. B) Probe test after training Schema B, confirming part A. C) Performance over 7 trials of retraining on Schema A. Both groups retained Schema A, though the control group recovered from a very minimal forgetting of the schema initially. D) Probe test after retraining on Schema A, confirming part C.

Figure 7 shows the weights of the network in the control condition after training on the third experiment. Weights from the context pattern to the mPFC now show two distinct schemas, with stronger patterns than seen in the first experiment. Weights from mPFC to the AC show two sets of neurons created by gating. Weights to the dHPC now show twice the amount of triplets as before, reflecting that triplets from two schemas are being encoded.

**Figure 7.**
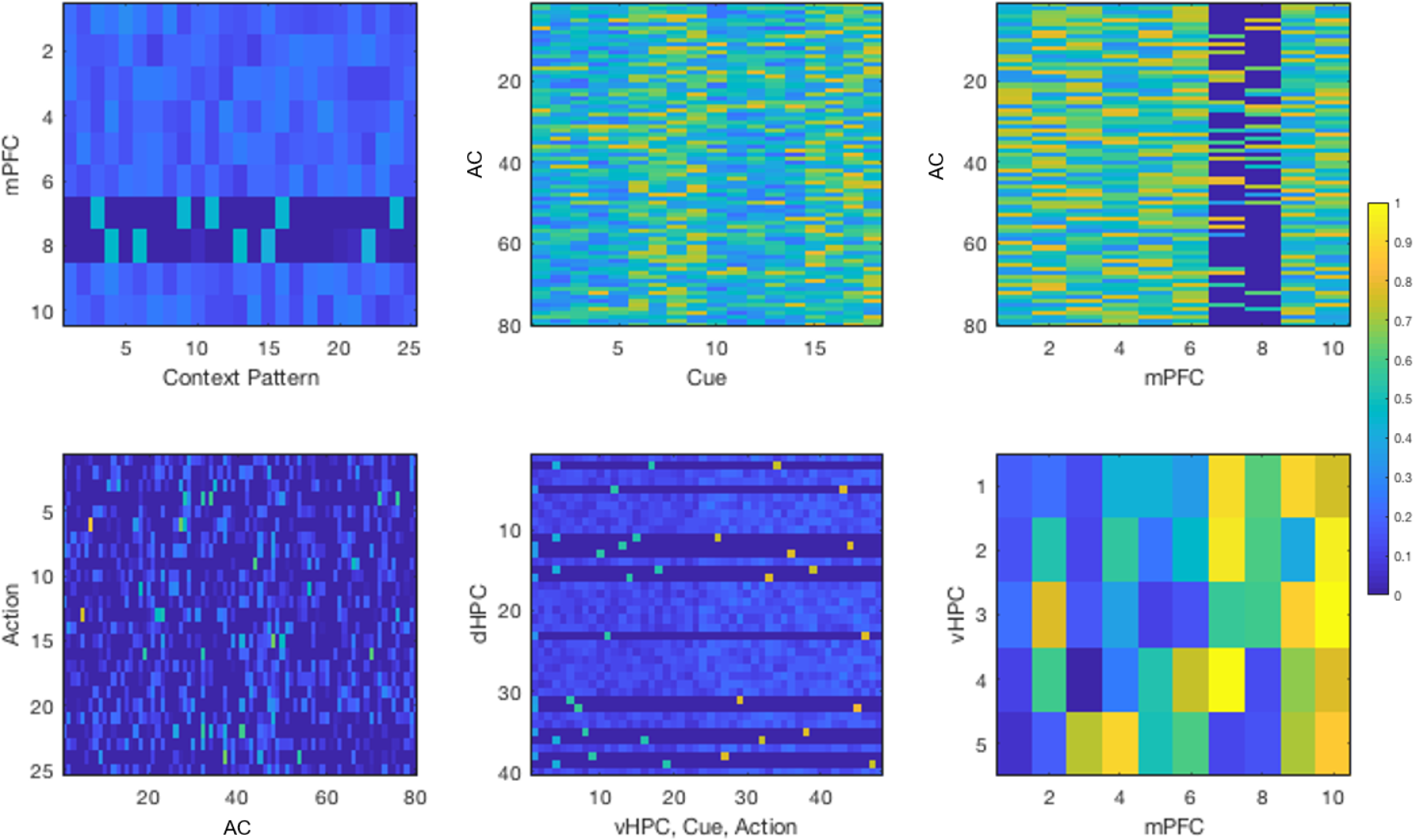
Weight matrices of the control network after simulating all of the second experiment. Rows represent post-synaptic layers and columns represent presynaptic layers.

The results of the second experiment show that our network is able to learn multiple schemas in succession, without forgetting prior schemas. We were thus able to match the effect seen in Tse et al. (2007). A small difference is that our network retained information about Schema A much better than their experiments for both of their HPC-lesioned and control groups.

### Experiment 3

The aim of the third experiment was to show the necessity of the schema in learning, as opposed to the agent simply getting better at learning with time. This was done by showing that a consistent schema is easier to learn and maintain than an inconsistent schema, as seen in the results in Figure 8A. This occurred in our model because an overlapping set of AC neurons was gated in all the inconsistent condition of Schema B and scrambled Schema B. This led to a confusion of the network weights as it was attempting to learn two layouts. This is confirmed in the probe tests performed after training, in Figure 8B. When adding new PAs to the consistent and inconsistent schemas in Figure 8C, the model was able to learn the new information better in the consistent schema.

**Figure 8.**
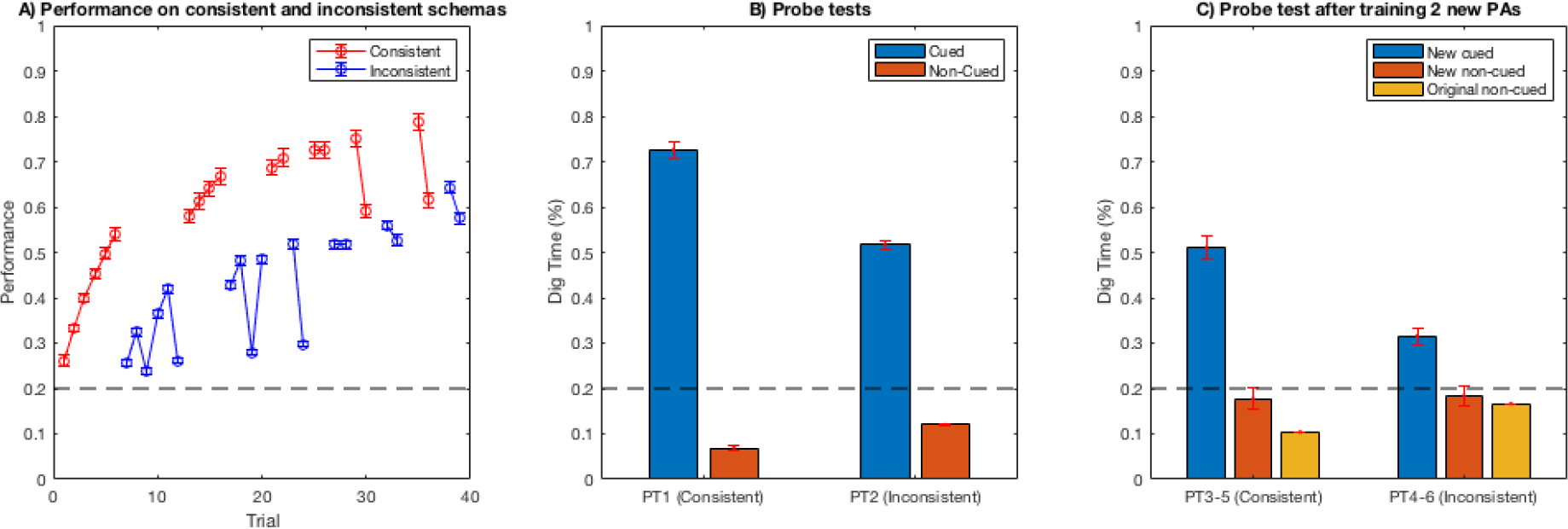
Results of replicating the third experiment of Tse et al. A) The performance over 40 trials of training, alternating between consistent and inconsistent schemas. While both conditions were able to learn, the inconsistent schema had worse performance. B) Probe tests of the consistent and inconsistent schemas show better performance of the consistent condition. C) Probe tests after one day of training two new PAs in both the consistent and inconsistent schemas. The new PAs were learned with greater performance in the consistent schema.

Figure 9 shows the weights of the network after training on the third experiment. The weights look very similar to the second experiment, as both encode two distinct schemas. The inconsistency of Schema B is not visible in the weights, as Schema B and scrambled Schema B have the same well locations. Only the spatial locations of the wells, not the flavors, are input to the context pattern.

**Figure 9.**
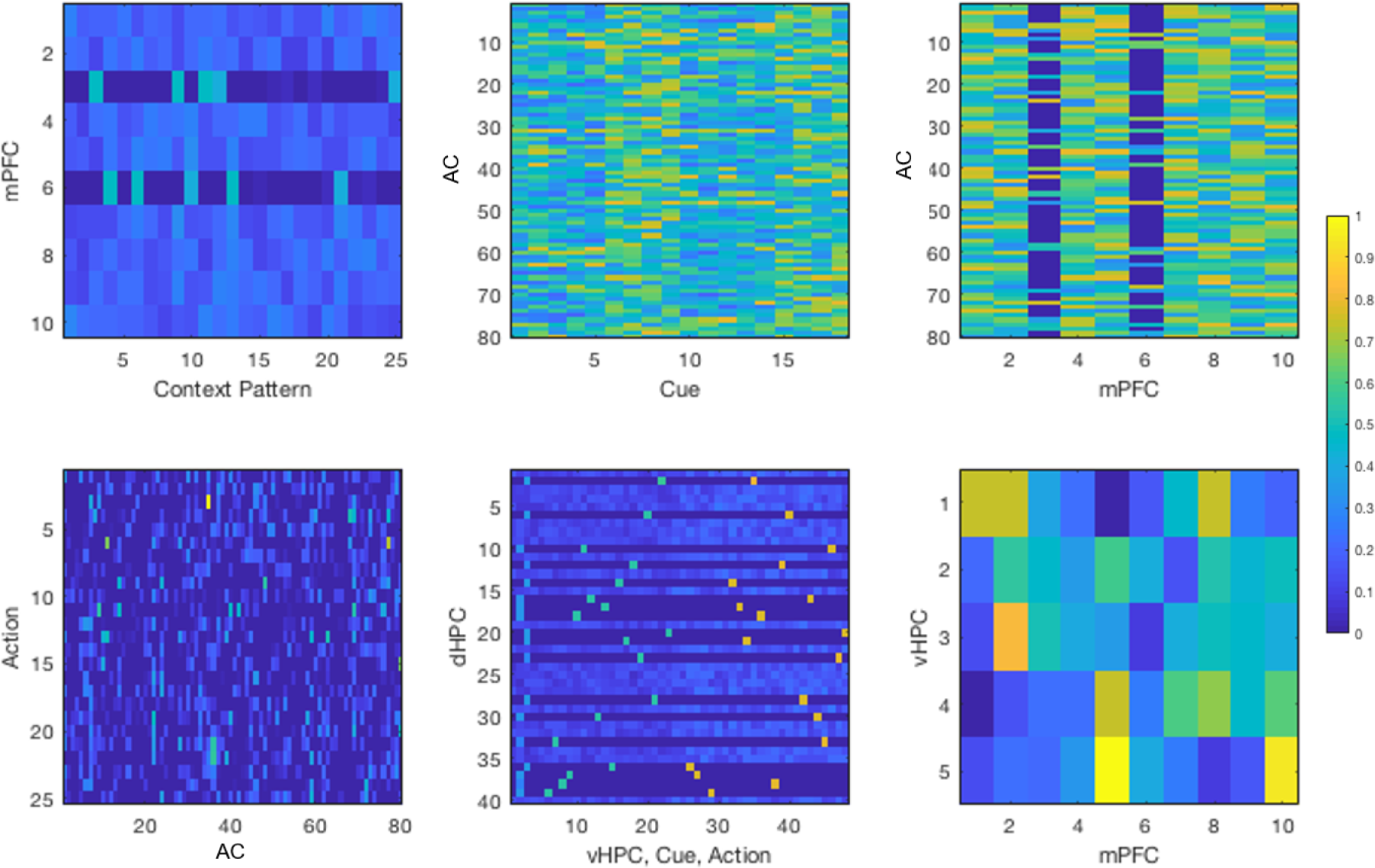
Weight matrices of the network after simulating all of the third experiment. Rows represent post-synaptic layers and columns represent presynaptic layers.

In our third experiment, we showed that learning new information within a consistent schema has better performance than an inconsistent schema. However, a large difference in Tse et al. (2007) was that there were almost no signs of learning in the probe tests for the inconsistent condition, suggesting that there may be an additional mechanism not addressed in our model that only develops schema representations when they are seen consistently.

### Neural Activity

To gain a better understanding of the network activity, we plotted the neuron traces of the mPFC, vHPC, and dHPC before winner-take-all was applied. Figure 10A shows the neuron activities for the first two experiments, with Schema A and new PAs for Experiment 1, and Schema B with a retraining of Schema A for Experiment 2. Each colored line in the figure represents the activity of a single neuron in a single simulated rat. The black vertical lines separate epochs into trials. The neuron traces show that the mPFC chooses a single neuron to represent Schema A and continues to strengthen its weights as it is trained. When new PAs are introduced, the same neuron is still active, but decreases in activity. When Schema B is introduced at the beginning of Experiment 1, a new neuron is chosen, and when Schema A is retrained immediately after training Schema B, the original winning neuron returns. By viewing the spacing between black vertical lines, we see that the large spacing for trials with new PAs indicates that more epochs of training occur during those times. This is due to the novelty and familiarity detection from the neuromodulator. vHPC activity in Figure 10B shows the same effect, except that the weights of all of the neurons together increase and decrease, as they are all affected by the rising and falling activity levels of mPFC. dHPC works similarly, as seen in Figure 10, although different winners are chosen at every epoch. For the dHPC we display neural traces for the first 100 epochs in the first trial epochs, due to the frequent switching of winners. Experiment 3 generates similar activity patterns to the other experiments (Figure 11).

**Figure 10.**
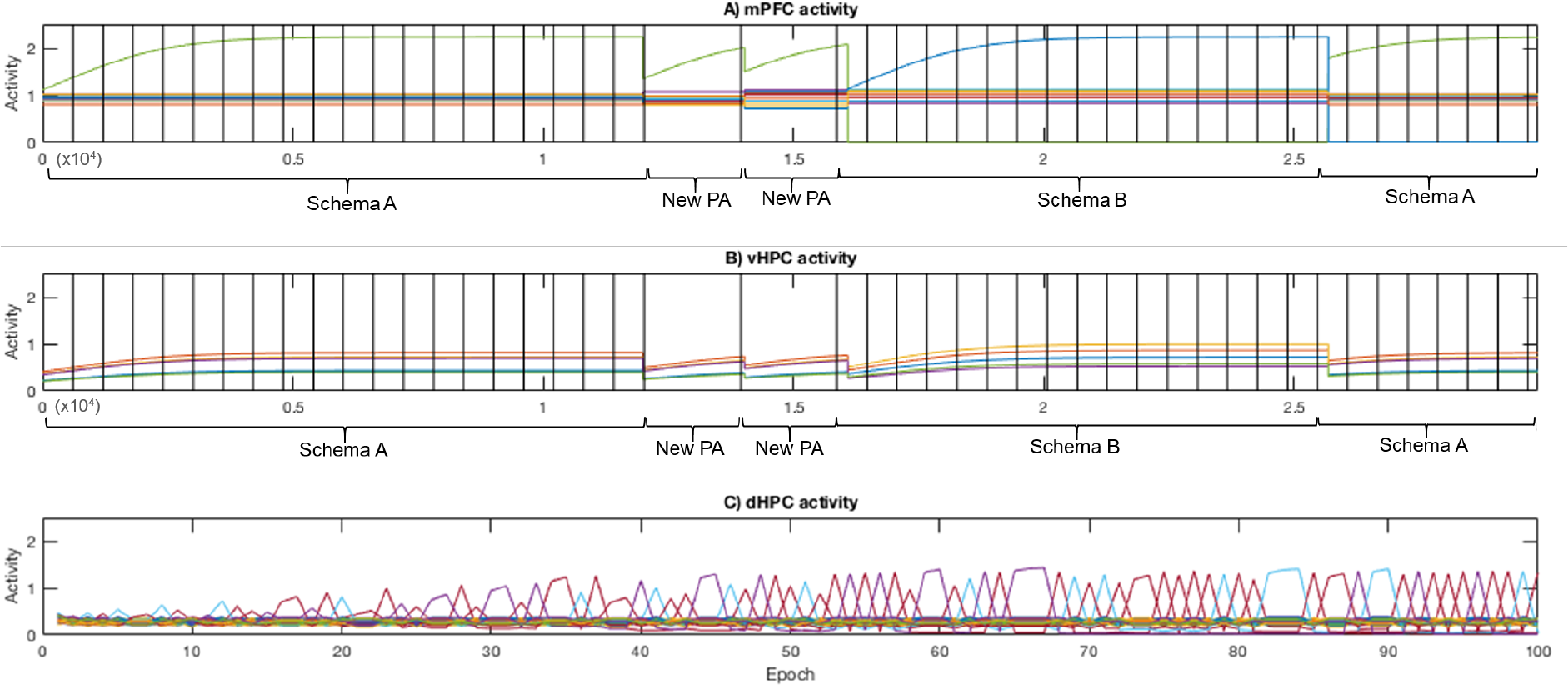
Neural activity for the network of a single simulated rat while performing the first and second experiments combined. Each colored line represents the activity of one neuron. Each vertical black line represents the separation of epochs into trials. Activity is measured before winner-take-all is applied. A) The mPFC activities for the first and second experiments are shown in sequence, with the training of Schema A and two instances of new PAs for the first experiment, and the training of Schema B and return of Schema A for the second experiment. When training on Schema A, one mPFC neuron is consistently chosen as winner, as its weight values increase over time. When new PAs are introduced, the winner remains the same, but decreases in activity. At the start of Experiment 2, Schema B is introduced and a new mPFC neuron wins. The number of epochs greatly increases due to the novelty. When Schema A is retrained, the original winning neuron returns. B) The neurons in vHPC follow the same general trend as the mPFC neurons. However, all neurons increase and decrease together, as they all take input from the winning mPFC neuron. C) Activity of the dHPC neurons for the first 100 epochs of the first trial is shown. As the density of switching of dHPC winners occurs at every epoch rather than at every trial, it is necessary to display the activity at the epoch level. At each epoch, a different dHPC neuron is selected to have its weights increase, as a different triplet is present each time. The activities of winning neurons gradually increase for 40 epochs, remaining stable afterwards.

**Figure 11.**
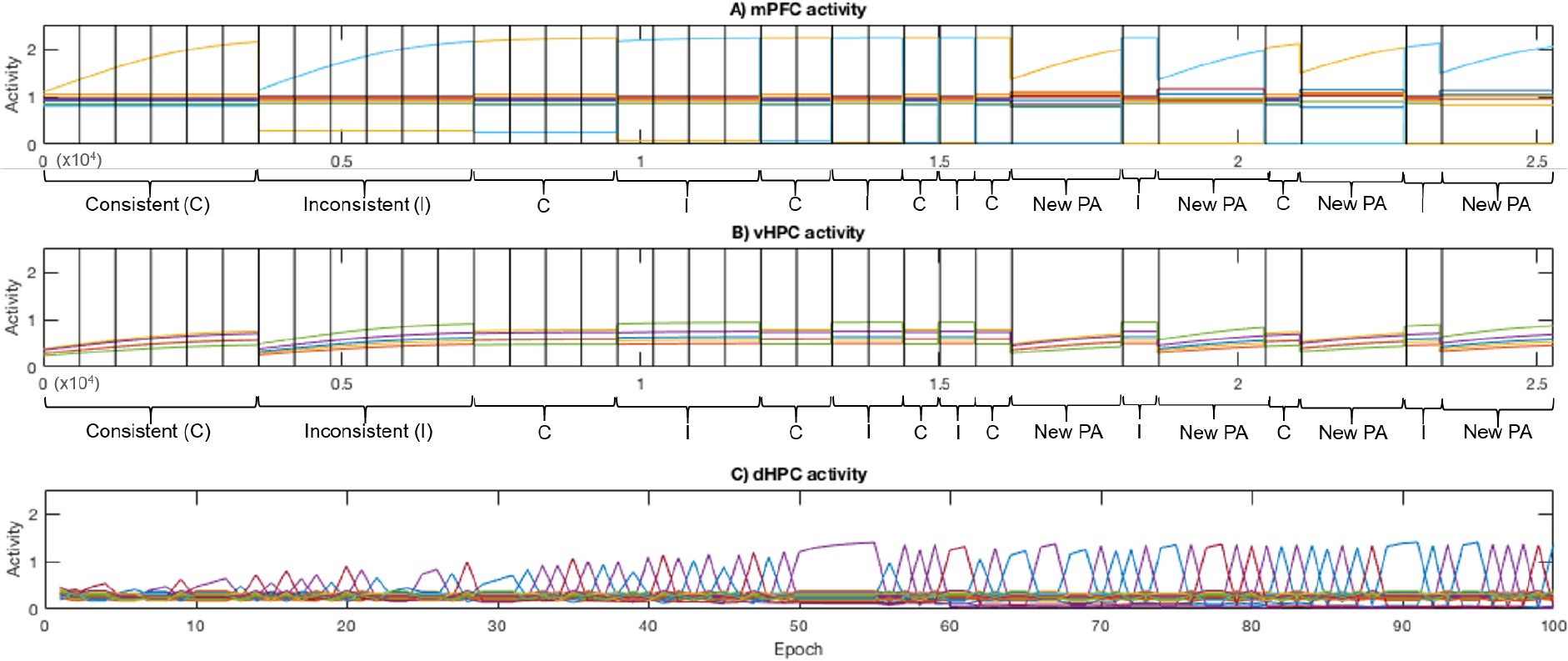
Neural activity for a single network while performing the third experiment. A) mPFC activity shows that one neuron’s weights increased in the consistent condition and that another neuron’s weights increased for the inconsistent condition. B) vHPC activity behaves similarly to Experiments 1 and 2. C) dHPC activity behaves similarly to Experiments 1 and 2.

To show the effects of schema familiarity and novelty on neuromodulator activity, Figure 12 displays the activities of the familiarity module, novelty module, and neuromodulator over the first 21 trials of Experiment 1. Due to the Hebbian learning of connections from the mPFC to the familiarity module, familiarity increases over time. Since it uses a sigmoidal transfer function, the increase is step-like, increasing from 0 to 1 swiftly. Novelty starts high when Schema A is introduced, and quickly drops to 0 due to the anti-Hebbian learning rule. When two new PAs are introduced, novelty returns to a high state. Taking the product of novelty and familiarity, the activity of the neuromodulator increases only when familiarity and novelty are high. Since the activity of the neuromodulator is proportional to the number of training epochs in a trial, this leads to the desired behavior of increased learning when new PAs are introduced to an existing schema.

**Figure 12.**
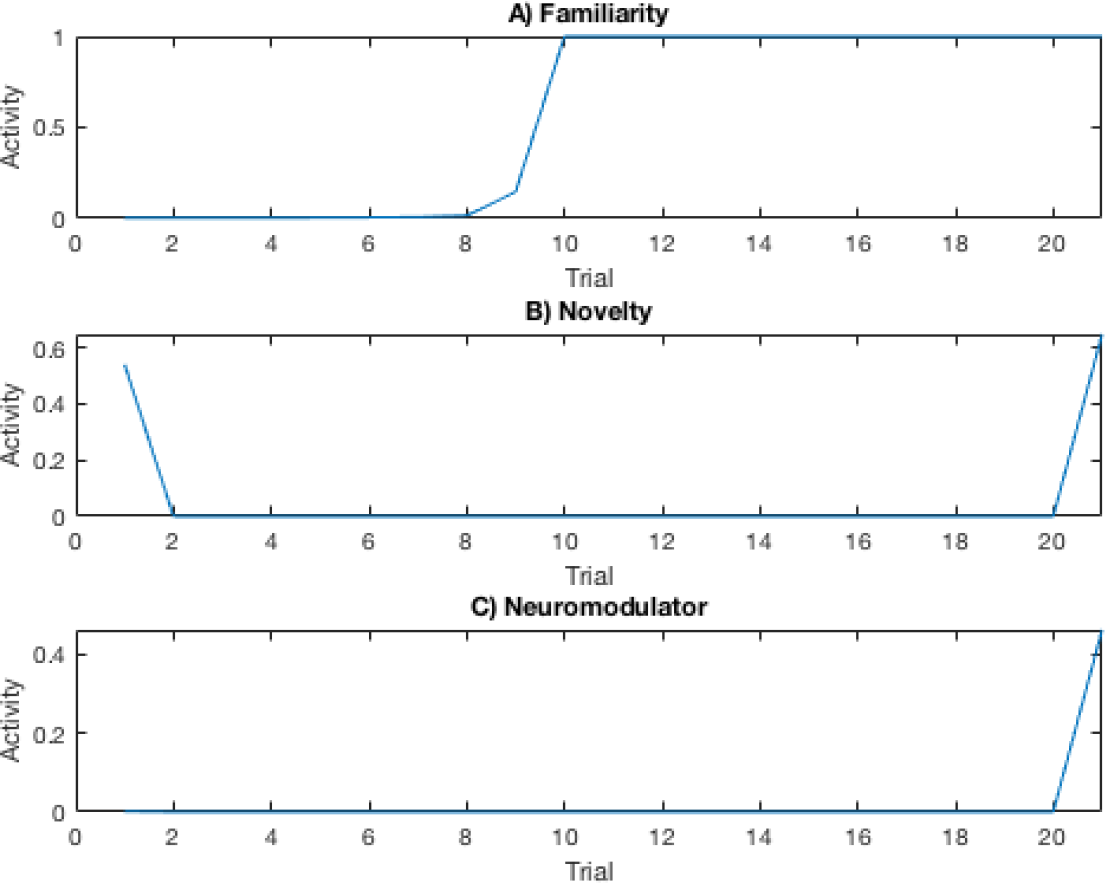
Combining familiarity and novelty for neuromodulation. A) Familiarity increases as the network is trained on a schema. Due to the sigmoidal transfer function, familiarity activity is step-like, going from unfamiliar to familiar over some training. B) Novelty starts at a high value whenever there are new PAs, which occurs on the first trial when the schema is introduced, and on the 21st trial when two PAs are replaced. C) At each trial, the product of familiarity and novelty lead to an increase of activity when schema familiarity is high and novelty is high.

We also tracked the activity of the neuromodulator for all three experiments. Figure 13A shows the number of training epochs in each trial of the first and second experiments. As a reminder, each epoch within a trial consists of an Indexing Phase, a Free Phase, and a Clamped Phase. The first 21 trials of the first experiment are on the left of the black vertical line, and the remaining trials to the right of the vertical line are for the second experiment. Each individual colored line represents the neuromodulator activity of a single run, with 20 runs total. When new PAs were introduced on trial 21 of the first experiment, the familiarity of Schema A multiplied by the novelty of the new stimuli caused a sharp spike in neuromodulator activity, increasing the number of epochs for those trials. The neuromodulator activity for the second experiment remained low, as there were no new PAs introduced. Figure 13 shows the index of the winning mPFC neuron in each trial, with a different colored line for each individual run. Every time the schema changes, the index of the winning neuron changes accordingly. Figures 13C and 13D show similar effects, with large increases in neuromodulator activity when there is a new PA and switching of active schema neurons whenever the schema changes from consistent to inconsistent and vice versa. The white regions represent consistent schema trials while the gray shaded regions represent inconsistent schema trials.

**Figure 13.**
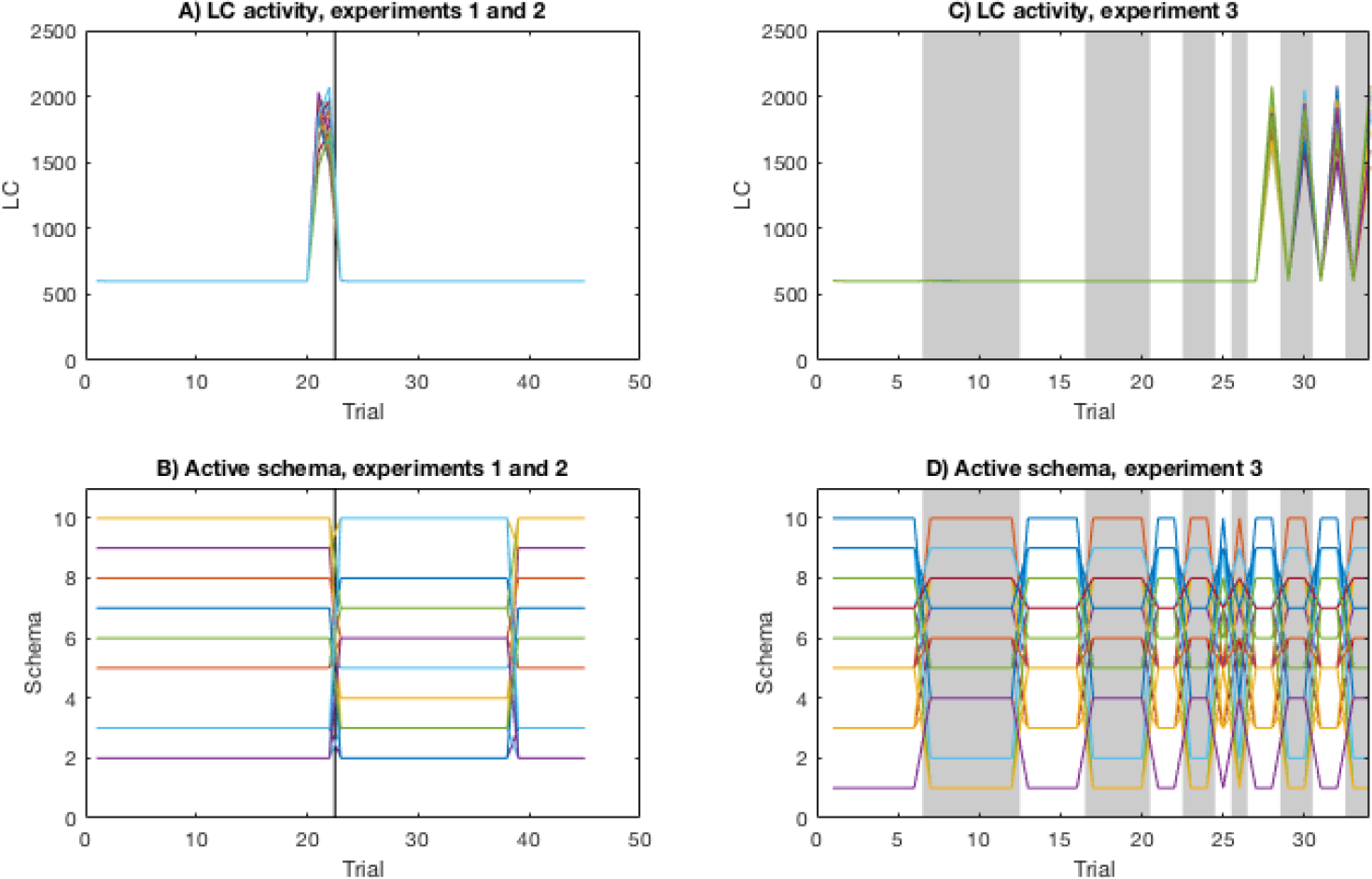
Neuromodulator activity and winning schemas. Each colored line represents the activity of the neuromodulator from an individual network out of 20. A) Neuromodulator activity expressed by the number of epochs trained per trial. The vertical black line separates experiments 1 and 2. On trial 21, the number of epochs rises sharply due to the combination of familiar schema and novel stimulus. B) Index of winning mPFC neuron in each trial of experiments 1 and 2. The index switches clearly each time the schema switches. C) Neuromodulator activity expressed by the number of epochs per trial for Experiment 3. Each time a new PA is introduced within a familiar schema, the number of epochs increases sharply. D) Index of winning mPFC neuron in each trial of Experiment 3.

### Effects of neuromodulation

To study how the neuromodulator influences learning, we removed connections to and from the novelty and familiarity modules in the network. Rather than boosting the number of epochs using neuromodulator activity, we used a constant number of epochs for each trial. We repeated the first part of Experiment 1 with the same number of epochs for every trial, trying different values for the constant number of epochs. As before, the network was trained on Schema A for 20 trials and training a new PA was introduced on the 21st trial. As seen in Figure 14, training on more epochs per trial leads to faster learning and better performance on the new PA. It is therefore not required to have a neuromodulator for rapid learning of new information. However, the number of total epochs over all trials can be greatly reduced if the model is able to detect novelty within the schema and adjust the learning accordingly. Compared to the conditions with a flat number of epochs per trial, we found that our original network with the neuromodulator had better performance on the new PAs than the other successful conditions, despite having fewer total epochs. It was therefore able to conserve network training time overall.

**Figure 14.**
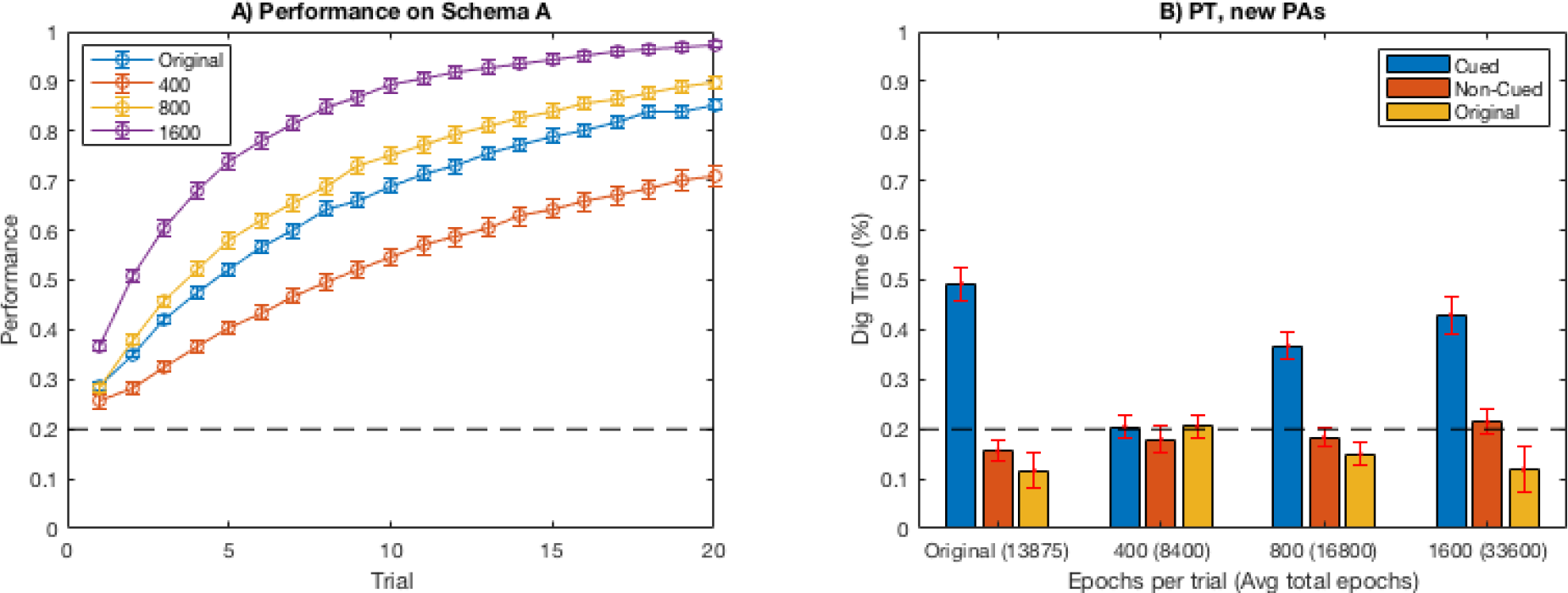
A) Performances of each condition. The blue line represents the original network and the remaining lines use a flat number of epochs as indicated in the legend. All conditions are able to learn the schema, but with different learning rates. B) Probe tests of each condition after introducing a new PA. The average number of epochs for each trial is displayed in parentheses. The performance of the probe test increases as more epochs are trained per trial. However, the original network can get a performance equivalent to the other conditions, but with far fewer epochs of training.

### Sparsity

An idea following from Tse et al. (2007) is that separation of information into schemas can be used to prevent catastrophic forgetting. In our network, this is highly dependent on the effects of the sparsity of the weight matrix between the vHPC and the AC layer. To test sparsity, we trained two schemas, A and B, in succession. During training, we varied the parameter P, which was the proportion of weights from vHPC to the AC layer that were set to zero. The results can be seen in Figure 15. For small values of P, the weights are so sparse that there are too few AC neurons to learn the task. For large values of P, the multimodal neurons selected for each schema overlaps such that Schema A is forgotten when Schema B is introduced. At an intermediate value of P, an adequate number of AC neurons is assigned to each schema, with little overlap such that catastrophic forgetting is prevented.

**Figure 15.**
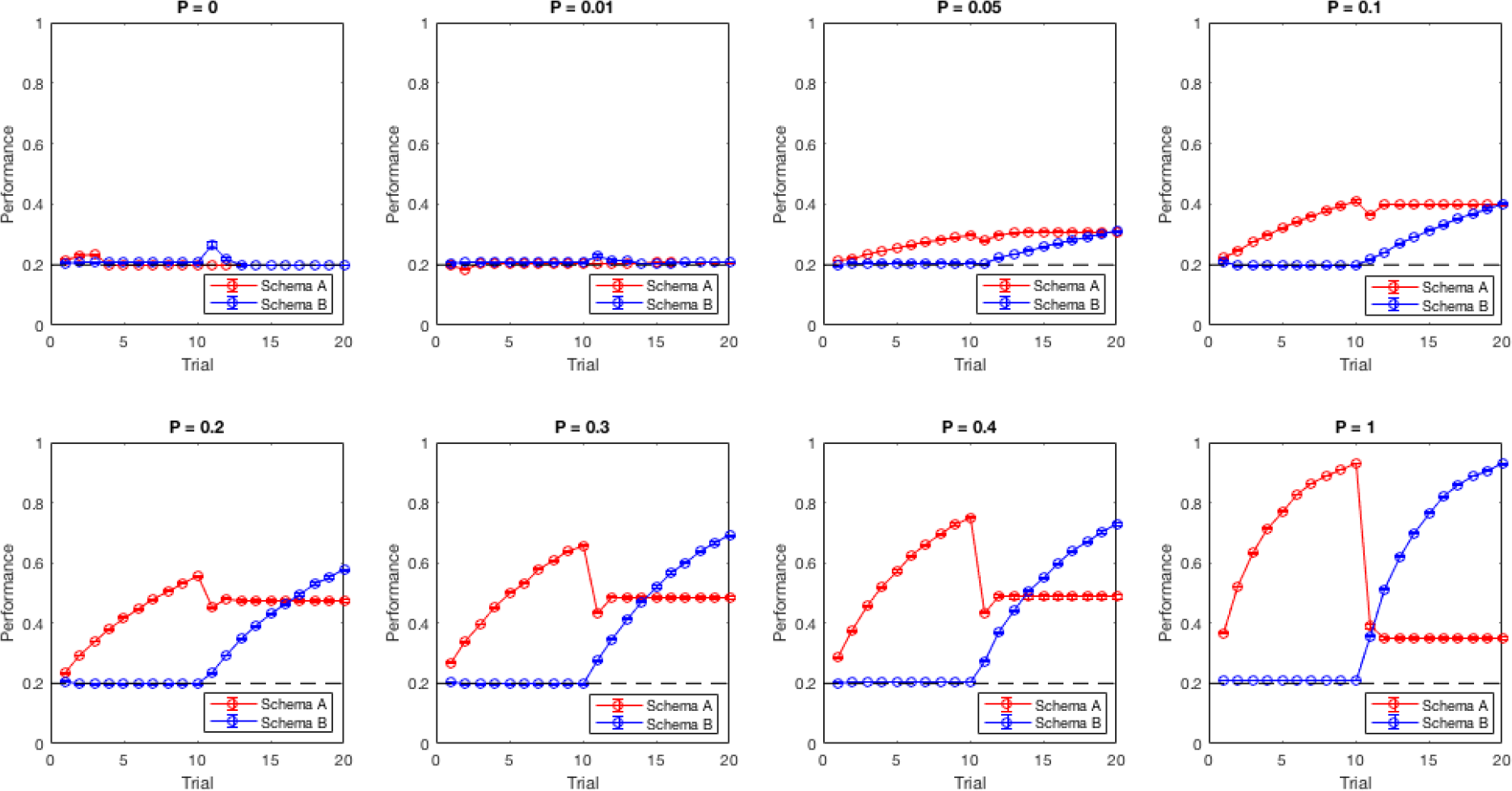
Performance of training Schema A for 10 trials and Schema B for 10 trials, each with a different P sparsity value. At low P values, no learning occurs. At high values, learning occurs but there is a steep drop in performance of Schema A when Schema B is introduced. At an intermediate level of sparsity, both schemas can be learned with an acceptably small amount of forgetting.

Since some of our results depended on the arrangement of the weight matrix for gating, it would be useful to see if there are different methods of sparsification that would yield better or worse results. While we sparsified the matrix through random patterns of positive weights, the matrix could be more structured so as to guarantee the avoidance of overlaps, or to coordinate with gating of other AC layers to support a hierarchical structure. As the sparsity level greatly impacted the performance of our network, this suggests that real nervous system connectivity makes a similar tradeoff between high sparsity, which leads to clean pattern separation, and low sparsity, which is required to adequately represent the features necessary for learning.

## Discussion

By replicating experiments from Tse et al. (2007), we showed that our biologically plausible neural network was able to learn schemas over time and quickly assimilate new information if it was consistent with a prior schema. The network was also able to learn multiple schemas without catastrophic forgetting, by maintaining separate sets of AC neurons for tasks within different schemas. Moreover, the consistency of the presentation of the schema affected how well new information could be assimilated into it.

The network highlights diverse roles of indexing by the HPC. Experimental literature shows that the spatial selectivity of place cells decreases along the dorsal-ventral axis of the HPC (Jung et al., 1994). Eichenbaum (2017) also proposes that specific memories are represented in the dHPC whereas contextual information is represented in the vHPC. Combined with indexing theory, our model shows that indexing may occur in hierarchical levels. For intermediate levels, the indexing separates representations of objects and tasks by the contexts in which they belong. This modularity makes it less likely that learning new information in new contexts will overwrite previously learned information in old contexts. The indexing of the dorsal HPC is necessary for driving the consolidation processes that transfer information to long term storage.

Our work is comparable to a recent approach to avoiding catastrophic forgetting by Nakano and Hattori (2017), in which the intermediate layers of a deep neural network are gated by patterns that differ by context. A paper by Masse, Grant, and Freedman (2018) has the similar idea of using CHL as a plausible deep representation of information, and applying "pseudopatterns" alongside their regular training patterns for better separation. Our experiments explain in more depth how these patterns are formed and employed throughout different stages of the learning process. The central location of the hippocampus makes it a likely candidate for effecting context-dependent gating within the network. Its connections to the mPFC allow the formation of context-dependent indices, and its wide connectivity to the whole neocortex gives it the ability to gate information at many levels of the hierarchy.

There are differing opinions on how interaction between the mPFC and HPC is involved in context-dependent tasks. The SLIMM model suggests a competitive relationship, with activation of the mPFC inhibiting the HPC when a stimulus is congruent with a prior schema. On the other hand, Preston and Eichenbaum (2013) suggest a more cooperative interaction, with the mPFC drawing specific memories from the vHPC, and in turn influencing the dHPC via entorhinal inputs. As the mPFC is also known to mediate attention shifting in context-dependent tasks (Birrell & Brown, 2000), it is likely that shifts in schemas cause the mPFC to change the activity of the HPC. In our experiment, the presence of new PAs within an existing schema should require both the mPFC and HPC to express the familiarity of the schema and novelty of the new PAs. Our model shows that a top down control of the HPC by the mPFC is able to change which specific neurons were active in the HPC and effectively separate tasks by schema.

Moreover, the gating patterns are not employed evenly at all times, but depend heavily on external factors such as novelty, uncertainty, and reward, therefore, our model predicts that gating of context information is controlled by neuromodulatory areas. This could explain how the brain controls phases of learning in such a flexible manner. Our model suggests that schema familiarity could be detected by the mPFC and novelty could be detected by the hippocampus. We propose that neuromodulation monitors these signals and amplifies learning and consolidation of new (i.e. novel or unfamiliar) information while sparing old information. Our simulated neuromodulator may have biological correlates in the LC, as it reacts to sudden changes in schemas and causes changes in theta oscillations within the HPC. However, the combination of novelty and familiarity is also reminiscent of the BF functionality suggested by Yu and Dayan (2005), in that the level of uncertainty is framed within a specific context. Other areas such as the dopaminergic ventral tegmental area (VTA) may be involved in neuromodulatory gating as well, as it has inbound and outbound connections with the hippocampus that control learning according to reward and novelty (Otmakhova, Duzel, Deutch, & Lisman, 2013).

We predict that the outcomes of our model could be tested experimentally by deactivating the LC or BF and testing whether this impairs the time it takes to learn the Tse et al. (2007) task. We would also expect that severing connections from the mPFC to the HPC would cause catastrophic forgetting, as intermediate representations would not be properly gated. It may also be possible to lesion the HPC and artificially gate on brain areas storing intermediate representations to see if this prevents catastrophic forgetting.

In addition to the neurobiological implications of our model, our work could have practical applications to a range of tasks in Artificial Intelligence and robotics. In the future, we hope to test our model on a variety of other datasets, particularly those used in machine learning and robotics applications. For instance, rather than learning the locations of food, our model can also learn the general layout of objects in a household. Different schemas formed would then correspond to different rooms, which can aid various robotic behaviors such as retrieving objects and interacting with people. The increased complexity of the environment may require the addition of more multimodal association layers in the network, which would test the scalability of context-based gating. Rather than having just a vHPC and dHPC, the model would include multiple HPC layers along the dorsal-ventral axis, one for each of the layers in the representation stream. These layers would then index at different levels of spatial, temporal, or conceptual hierarchies, depending on the task at hand. The use of more layers presents the possibility of starting from raw visual input as opposed to labeled objects, for an entirely end-to-end approach to context-based task learning.

## Acknowledgments

We acknowledge the participants of the Telluride Neuromorphic Workshop 2017, including Xinyun Zou, Brent Komer, Georgios Detorakis, and Scott Koziol, who worked on a preliminary project leading to the creation of this model. Our work was funded in part by Toyota Motor North America.

## References

Abraham, W. C., & Robins, A. (2005). Memory retention–the synaptic stability versus plasticity dilemma. Trends in neurosciences, 28 (2), 73–78.

Aston-Jones, G., & Cohen, J. D. (2005). Adaptive gain and the role of the locus coeruleus–norepinephrine system in optimal performance. Journal of Comparative Neurology, 493 (1), 99–110.

Baldi, P., & Pineda, F. (1991). Contrastive learning and neural oscillations. Neural Computation, 3 (4), 526–545.

Baxter, M. G., & Chiba, A. A. (1999). Cognitive functions of the basal forebrain. Current opinion in neurobiology, 9 (2), 178–183.

Berridge, C. W., & Foote, S. L. (1991). Effects of locus coeruleus activation on electroencephalographic activity in neocortex and hippocampus. Journal of Neuroscience, 11 (10), 3135–3145.

Birrell, J. M., & Brown, V. J. (2000). Medial frontal cortex mediates perceptual attentional set shifting in the rat. Journal of Neuroscience, 20 (11), 4320–4324.

Diba, K., & Buzsáki, G. (2007). Forward and reverse hippocampal place-cell sequences during ripples. Nature neuroscience, 10 (10), 1241.

Eichenbaum, H. (2017). Prefrontal–hippocampal interactions in episodic memory. Nature Reviews Neuroscience, 18 (9), 547.

French, R. M. (1999). Catastrophic forgetting in connectionist networks. Trends in cognitive sciences, 3 (4), 128–135.

Hawkins, J., Ahmad, S., & Cui, Y. (2017). A theory of how columns in the neocortex enable learning the structure of the world. Frontiers in neural circuits, 11, 81.

Jung, M. W., Wiener, S. I., & McNaughton, B. L. (1994). Comparison of spatial firing characteristics of units in dorsal and ventral hippocampus of the rat. Journal of Neuroscience, 14 (12), 7347–7356.

Kirkpatrick, J., Pascanu, R., Rabinowitz, N., Veness, J., Desjardins, G., Rusu, A. A., … others (2017). Overcoming catastrophic forgetting in neural networks. Proceedings of the national academy of sciences, 201611835.

Kitamura, T., Ogawa, S. K., Roy, D. S., Okuyama, T., Morrissey, M. D., Smith, L. M., … Tonegawa, S. (2017). Engrams and circuits crucial for systems consolidation of a memory. Science, 356 (6333), 73–78.

Krichmar, J. L. (2008). The neuromodulatory system: a framework for survival and adaptive behavior in a challenging world. Adaptive Behavior, 16 (6), 385–399.

Kumaran, D., Hassabis, D., & McClelland, J. L. (2016). What learning systems do intelligent agents need? complementary learning systems theory updated. Trends in cognitive sciences, 20 (7), 512–534.

LeCun, Y., Bengio, Y., & Hinton, G. (2015). Deep learning. nature, 521 (7553), 436.

Masse, N. Y., Grant, G. D., & Freedman, D. J. (2018). Alleviating catastrophic forgetting using context-dependent gating and synaptic stabilization. arXiv preprint arXiv:1802.01569.

McClelland, J. L., McNaughton, B. L., & O’Reilly, R. C. (1995). Why there are complementary learning systems in the hippocampus and neocortex: insights from the successes and failures of connectionist models of learning and memory. Psychological review, 102 (3), 419.

Mermillod, M., Bugaiska, A., & Bonin, P. (2013). The stability-plasticity dilemma: Investigating the continuum from catastrophic forgetting to age-limited learning effects. Frontiers in psychology, 4, 504.

Movellan, J. R. (1991). Contrastive Hebbian learning in the continuous hopfield model. In Connectionist models (pp. 10–17). Elsevier.

Nakano, S., & Hattori, M. (2017). Reduction of catastrophic forgetting in multilayer neural networks trained by contrastive Hebbian learning with pseudorehearsal. In Computational intelligence and applications (IWCIA), 2017 IEEE 10th international workshop on (pp. 91–95).

Otmakhova, N., Duzel, E., Deutch, A. Y., & Lisman, J. (2013). The hippocampal-vta loop: the role of novelty and motivation in controlling the entry of information into long-term memory. In Intrinsically motivated learning in natural and artificial systems (pp. 235–254). Springer.

Preston, A. R., & Eichenbaum, H. (2013). Interplay of hippocampus and prefrontal cortex in memory. Current Biology, 23 (17), R764–R773.

Rumelhart, D. E., Hinton, G. E., & Williams, R. J. (1986). Learning representations by back-propagating errors. Nature, 323 (6088), 533.

Soltoggio, A., Stanley, K. O., & Risi, S. (2017). Born to learn: the inspiration, progress, and future of evolved plastic artificial neural networks. arXiv preprint arXiv:1703.10371.

Teyler, T. J., & DiScenna, P. (1986). The hippocampal memory indexing theory. Behavioral neuroscience, 100 (2), 147.

Tse, D., Langston, R. F., Kakeyama, M., Bethus, I., Spooner, P. A., Wood, E. R., … Morris, R. G. (2007). Schemas and memory consolidation. Science, 316 (5821), 76–82.

Tse, D., Takeuchi, T., Kakeyama, M., Kajii, Y., Okuno, H., Tohyama, C., … Morris, R. G. (2011). Schema-dependent gene activation and memory encoding in neocortex. Science, 333 (6044), 891–895.

Van Kesteren, M. T., Beul, S. F., Takashima, A., Henson, R. N., Ruiter, D. J., & Fernández, G. (2013). Differential roles for medial prefrontal and medial temporal cortices in schema-dependent encoding: from congruent to incongruent. Neuropsychologia, 51 (12), 2352–2359.

Wagatsuma, A., Okuyama, T., Sun, C., Smith, L. M., Abe, K., & Tonegawa, S. (2018). Locus coeruleus input to hippocampal CA3 drives single-trial learning of a novel context. Proceedings of the National Academy of Sciences, 115 (2), E310–E316.

Walling, S. G., Brown, R. A., Milway, J. S., Earle, A. G., & Harley, C. W. (2011). Selective tuning of hippocampal oscillations by phasic locus coeruleus activation in awake male rats. Hippocampus, 21 (11), 1250–1262.

Xie, X., & Seung, H. S. (2003). Equivalence of backpropagation and contrastive Hebbian learning in a layered network. Neural computation, 15 (2), 441–454.

Yu, A., & Dayan, P. (2005). Uncertainty, neuromodulation, and attention. Neuron, 46 (4), 681–692.

